# Evidence Scaling for Zero-Shot Protein Reasoning with Large Language Models

**DOI:** 10.64898/2026.09.27.754822

**Authors:** Zitong Hao, Chaoyang Wang, Dongyuan Li, Yiquan Wang

## Abstract

Large language models (LLMs) show emerging zero-shot capability for protein variant prediction, yet still lag behind specialized protein models. We ask whether this gap can be reduced by scaling access to biological evidence rather than adapting model parameters. We introduce BioEvidence, a training-free and model-agnostic interface that converts structural and evolutionary information from standard biological tools into compact evidence for frozen LLMs. On the ProteinGym benchmark, we observe evidence scaling: performance improves as evidence becomes richer. Structural and evolutionary evidence each improve performance, and combining them yields further gains, while mismatching the same evidence to the wrong variants degrades performance below the no-evidence baseline. Notably, BioEvidence enables zero-shot ranking to reach strong specialized protein predictors on matched evaluations, and the improvement persists on post-cutoff data released after the model’s knowledge cutoff. Evidence also interacts with conventional scaling: for GPT-5.6 Sol, evidence at low reasoning effort outperforms the no-evidence condition at medium effort, while a six-model analysis associates stronger no-evidence performance with larger margins over evolutionary rank fusion. These results identify external evidence as a complementary scaling axis for scientific prediction alongside model capability and inference effort.

## 1 Introduction

General-purpose LLMs can rank protein variants from a wild-type sequence, assay description, and candidate sequences, but still trail specialized predictors even with additional test-time compute (Arora et al., 2026). Unlike many specialists, they receive neither multiple-sequence alignments (MSAs) nor protein structures.

The comparison therefore conflates specialization with information access: GEMME (Laine et al., 2019) uses alignments, whereas ESM-IF (Hsu et al., 2022) conditions on structure. This motivates a question distinct from parameter adaptation: *how much can a frozen LLM improve when given explicit biological evidence?*

BioEvidence renders standard structural and evolutionary annotations as compact candidate-level evidence (Figure 1), without fine-tuning or learned adapters. Precomputation lets us vary evidence coverage and candidate correspondence while holding the model and task fixed.

**Figure 1.**
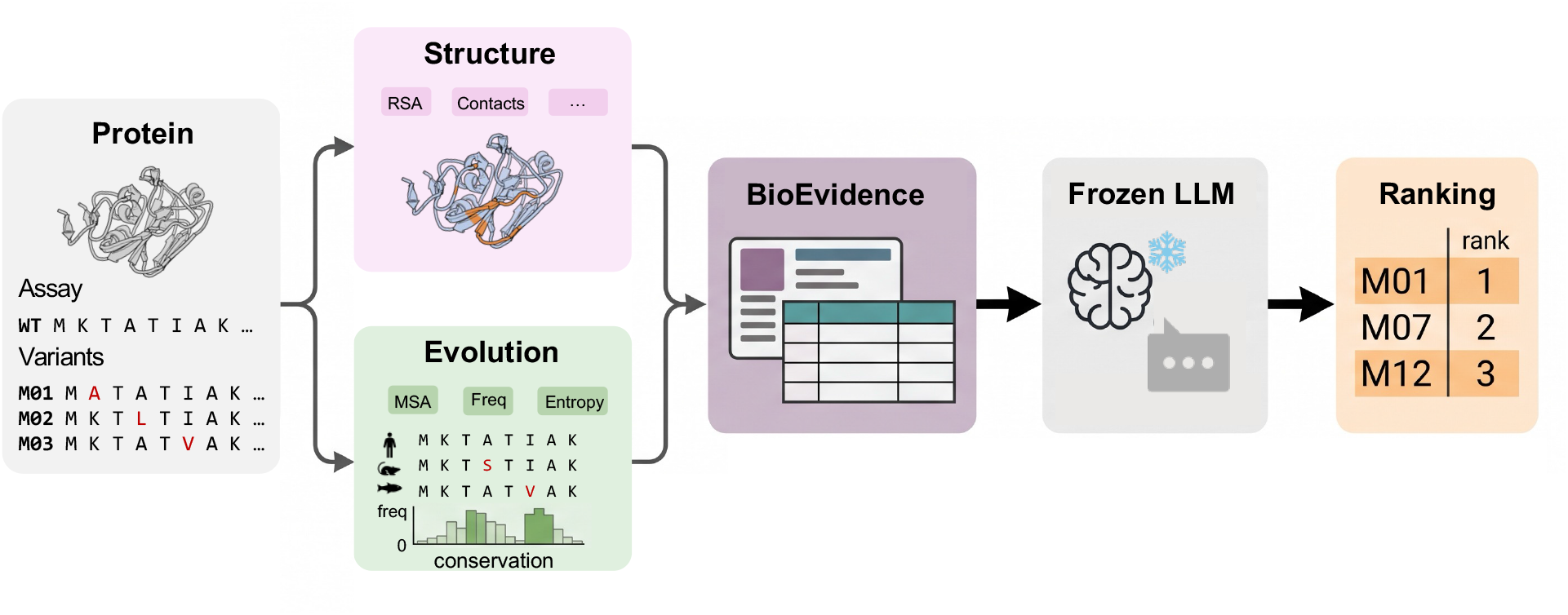
BioEvidence pipeline. Structural features and evolutionary statistics are computed for the candidate variants and rendered as compact evidence blocks for a frozen general-purpose LLM. The model parameters remain unchanged; only the biological evidence available at inference time varies.

We study evidence alongside model and inference scaling (Kaplan et al., 2020; Hoffmann et al., 2022; Wei et al., 2022; Snell et al., 2024): we vary coverage, shuffle candidate assignments, compare with specialists and post-cutoff assays, and test substitution for reasoning effort. A mechanical fusion of no-evidence predictions and an all-site evolutionary readout tests whether evidence prompting outperforms this fixed combination.

### Contributions

1. **Evidence scales performance**. For GPT-5.6 Sol, performance rises monotonically from 0.322 to 0.433 as coverage increases from 0% to 100%. Structural and evolutionary evidence help separately and combine best; mismatching evidence to variants instead reduces performance by 0.088.
2. **Evidence narrows the specialist gap**. Claude Opus 5 ranks above 91 of 95 published predictors on available coverage but remains 0.048 below VenusREM on matched candidates. GPT-6 Astra reaches similar matched performance to two leading specialists on its incomplete panel. A 25-assay post-cutoff evaluation provides preliminary evidence of transfer.
3. **Evidence can substitute for reasoning effort**. GPT-5.6 Sol at low effort with evidence exceeds medium effort without evidence by 0.104; the corresponding Opus 4.8 contrast is less evident.
4. **Gains beyond evolutionary fusion vary across models**. The advantage of evidence prompting over evolutionary rank fusion is associated with no-evidence performance (Pearson +0.789; Spearman +0.829; six models). This exploratory result does not identify an internal reasoning mechanism.

## 2 Related Work

### LLMs for protein prediction

We adopt the fixed ProteinGym draws and ranking protocol of PG-LLM (Arora et al., 2026; Notin et al., 2023a). Other approaches connect protein representations to language models: Prot2Chat (Wang et al., 2025) combines a ProteinMPNN (Dauparas et al., 2022) encoder with adaptation of the language model; ProTrek (Su et al., 2026) aligns sequence, structure, and functional text; and Cuttlefish (Jing et al., 2026) adapts structural tokens for LLM reasoning. STELLA (Jin et al., 2025) instead enables models to call scientific tools. Our study evaluates precomputed evidence supplied to an unmodified LLM, isolating evidence availability from learned interfaces and tool-selection policies.

### Specialized variant-effect predictors

Protein predictors differ in their inference inputs: sequence models such as ESM-1v (Meier et al., 2021); alignment-based methods such as GEMME (Laine et al., 2019); and structure-aware methods such as ESM-IF (Hsu et al., 2022). VenusREM (Tan et al., 2025) combines evolutionary and structural context and leads our main-benchmark reference set. We evaluate a text interface to related annotations, without adapting model parameters. Appendix D provides a broader comparison, including methods with different supervision and prediction targets.

### Evidence-grounded reasoning

In-context learning (Brown et al., 2020) and retrieval-augmented generation (Lewis et al., 2020) demonstrate how external context can influence LLM outputs. ReAct (Yao et al., 2023) and Toolformer (Schick et al., 2023) incorporate tool use, while ChemCrow (M. Bran et al., 2024) and Coscientist (Boiko et al., 2023) apply tools to scientific tasks. Here, evidence acquisition is fixed: the model neither retrieves information nor selects tools, making coverage and candidate correspondence controlled variables.

## 3 BIOEVIDENCE: Evidence for Frozen Language Models

### 3.1 ProteinGym Ranking Task

We evaluate on the ProteinGym substitution benchmark (Notin et al., 2023a) under the episode protocol of the PG-LLM suite (Arora et al., 2026): 217 assays, three fixed candidate draws per assay, 50 candidate variants per draw, giving 651 episodes. Each episode presents the wild-type sequence, a natural-language description of the assay, and the 50 candidates in a fixed randomized order; the model returns a ranking. These fixed draws support comparisons on identical candidates wherever episode-level reference predictions are available.

### 3.2 Biological Evidence Interface

BioEvidence attaches to each candidate a compact block assembled from standard tool outputs: secondary structure and relative solvent accessibility from DSSP (Kabsch & Sander, 1983) via Biopython (Cock et al., 2009), read off AlphaFold (Jumper et al., 2021) models retrieved from AlphaFold DB (Varadi et al., 2022) and gated on per-residue pLDDT, together with weighted residue frequencies from the alignment. The model receives these precomputed annotations through five input conditions:

#### R (raw)

Wild-type sequence, assay description, and all 50 candidate sequences, without biological annotations. This is our internal no-evidence control.

#### S (structural evidence)

R plus, for each mutated site: DSSP secondary structure, relative solvent accessibility, the AlphaFold per-residue pLDDT, and the non-local C*α* neighbours within 8 Å (sequence separation *>* 2) with their distances and count. The block is not uniform across candidates: where pLDDT falls below 70 the other three fields are withheld as unreliable and only the pLDDT is shown, and residues DSSP does not cover are rendered as *not computed* rather than silently omitted.

#### E (evolutionary evidence)

R plus, for each mutated site: alignment depth (raw and effective), site coverage, the weighted wild-type and mutant residue frequencies, and the site entropy.

#### SE (structure + evolution)

R plus both blocks, for every candidate.

#### SH (shuffled)

The SE evidence blocks, correctly formatted and identical in length and composition, but with their payloads permuted among candidates having the same number of substituted sites. Candidate identifiers and true position headers remain fixed. The permutation is a deterministic derangement within each non-singleton group (seed 20260915); singleton groups necessarily remain unchanged. SH holds prompt length and surface form fixed while disrupting most evidence–variant correspondences.

For multi-site variants, S, E, and SE contain one position-specific sub-block for every substitution. The SH control preserves the length, formatting, and biological vocabulary of SE while disrupting candidate correspondence. Across 651 episodes, 72 candidates in 25 episodes belonged to singleton site-count groups and therefore retained their own payloads; this exception is recorded in the shuffle manifest. Comparing SE with SH tests whether correctly assigned evidence contributes beyond these shared prompt properties.

### 3.3 Controlled Evidence Scaling

We supply evidence to 0%, 25%, 50%, 75%, or 100% of candidates using nested subsets. For each episode, candidate identifiers are shuffled by a deterministic pseudorandom seed derived from the first 16 hexadecimal digits of the episode identifier’s SHA-256 digest. The first 12, 25, and 37 candidates in that order receive evidence at the three intermediate doses (integer counts are rounded down), making each lower-dose set a subset of the next. Because the episode identifier includes the draw, selection is independent across draws. Retained evidence blocks are byte-identical across doses; candidates without evidence receive an explicit missing-evidence notice. Increasing coverage replaces notices with evidence blocks. This also increases prompt length: the 75% condition is 19.9% longer than 25% across the corpus (median per-episode increase 14.2%). The dose series therefore does not isolate length effects; SH supplies the complementary control (Section 3.2).

## 4 Experimental Setup

### 4.1 Models

We evaluate both R and SE conditions for six models: GPT-5.6 Sol, GPT-5.5, Claude Opus 5, Claude Opus 4.8, Claude Opus 4.7, and Claude Haiku 4.5. A further model, GPT-6 Astra, has only an SE condition and appears in the specialist and post-cutoff comparisons; it is excluded from SE−R and fusion-baseline analyses. We report reasoning-effort settings explicitly and distinguish comparisons at the same effort from those across effort tiers. The in-house Claude Opus 5 R/SE pair crosses effort tiers and therefore is not used in the primary fusion panel. Instead, we reconstruct candidate-level Claude Opus 5 max-effort R rankings from the public PG-LLM traces (Arora et al., 2026). This matches effort but crosses access channels; the in-house medium-R result is retained only as an R-source sensitivity check in Appendix C. A requested Claude Opus 5.5 condition was excluded after a served-model audit detected model substitution (Appendix A.7).

### 4.2 Evaluation and Statistics

We use the benchmark’s nested-macro Spearman correlation, *ρ*. Each episode is scored against measured fitness. Within each candidate draw, we first average assays belonging to the same protein and functional category, then average proteins within each category, and finally average the five categories with equal weight. The headline score is the mean across the three draws. Fitness values are never compared across assays.

For pairwise comparisons, both conditions are rescored on their shared episodes. For experiments involving several conditions, we use the intersection across all conditions unless otherwise stated. We distinguish these matched estimates from *own-coverage* scores, which use every episode scored by each entry and serve only as descriptive summaries. Confidence intervals for the primary paired contrasts use 10,000 protein-level bootstrap resamples and percentile 95% intervals (Efron, 1979). Each sampled protein carries all its assays and draws, with separate protein–category cells retained throughout the nested aggregation. This preserves the point estimator while accounting for dependence within proteins. Exploratory analyses and exceptions to this procedure are identified where reported.

### 4.3 Post-Cutoff Evaluation

We evaluate 25 of 59 assays designated as post-cutoff, selected before scoring because both evidence types could be built without assay-specific repair. Following the temporal benchmark, we average assays within each (study group, draw) cell and weight observed cells equally. Matched intervals use 10,000 paired study-group bootstrap resamples. Alignments and structures are rebuilt under a frozen protocol (Appendix A.3). This reduces opportunities to memorize assay measurements but cannot establish that related proteins were absent from training data.

### 4.4 Coverage and Failure Handling

Refusals, context overflows, and unparsable responses are recorded as missing rankings and reported separately. We do not assign them a Spearman score of zero. Coverage differs across conditions, and missing episodes are not random; matched comparisons therefore estimate performance conditional on successful ranking by all compared conditions. Own-coverage leaderboard positions do not establish paired superiority.

#### Channel calibration

In-house remote runs use CLI channels, whereas published LLM baselines use native APIs. Matched-model calibration finds lower GPT scores through our channel, but no detectable offset for Claude Opus 4.8 (Appendix A.4). We use in-house R conditions for controlled contrasts and explicitly label comparisons that cross channels.

## 5 Results

### 5.1 Ranking Performance Increases with Evidence Coverage

#### Correct candidate correspondence matters

For GPT-5.6 Sol at medium effort, own-coverage nested-macro Spearman is 0.234 under SH, 0.321 under R, and 0.434 under SE (Table 1). On the 580 episodes shared by all three conditions, SE improves over R by 0.113 (95% CI [0.091, 0.134]), whereas SH reduces performance by 0.088 ([−0.106, −0.071]). Because SH preserves the evidence content and prompt length while changing its assignment, these results show that correct candidate correspondence is important; prompt length alone cannot explain the SE improvement.

**Table 1:** Three-arm contrast for GPT-5.6 Sol at medium effort. Scores use own coverage; differences and paired-bootstrap 95% CIs use 580 shared episodes.

| Condition | $\rho$ | $\Delta$ vs. R | 95% CI |
| --- | --- | --- | --- |
| SH (shuffled evidence) | 0.234 | −0.088 | [−0.106, −0.071] |
| R (no evidence) | 0.321 | — | — |
| SE (structure + evolution) | 0.434 | +0.113 | [+0.091, +0.134] |

#### Performance increases monotonically with coverage

Relative to R, gains are +0.041, +0.072, +0.088, and +0.112 at 25%, 50%, 75%, and 100% coverage (Figure 2a). The ordering is stable across three coverage restrictions (maximum drift 0.002), establishing monotonicity over the tested doses, not a linear scaling law.

**Figure 2.**
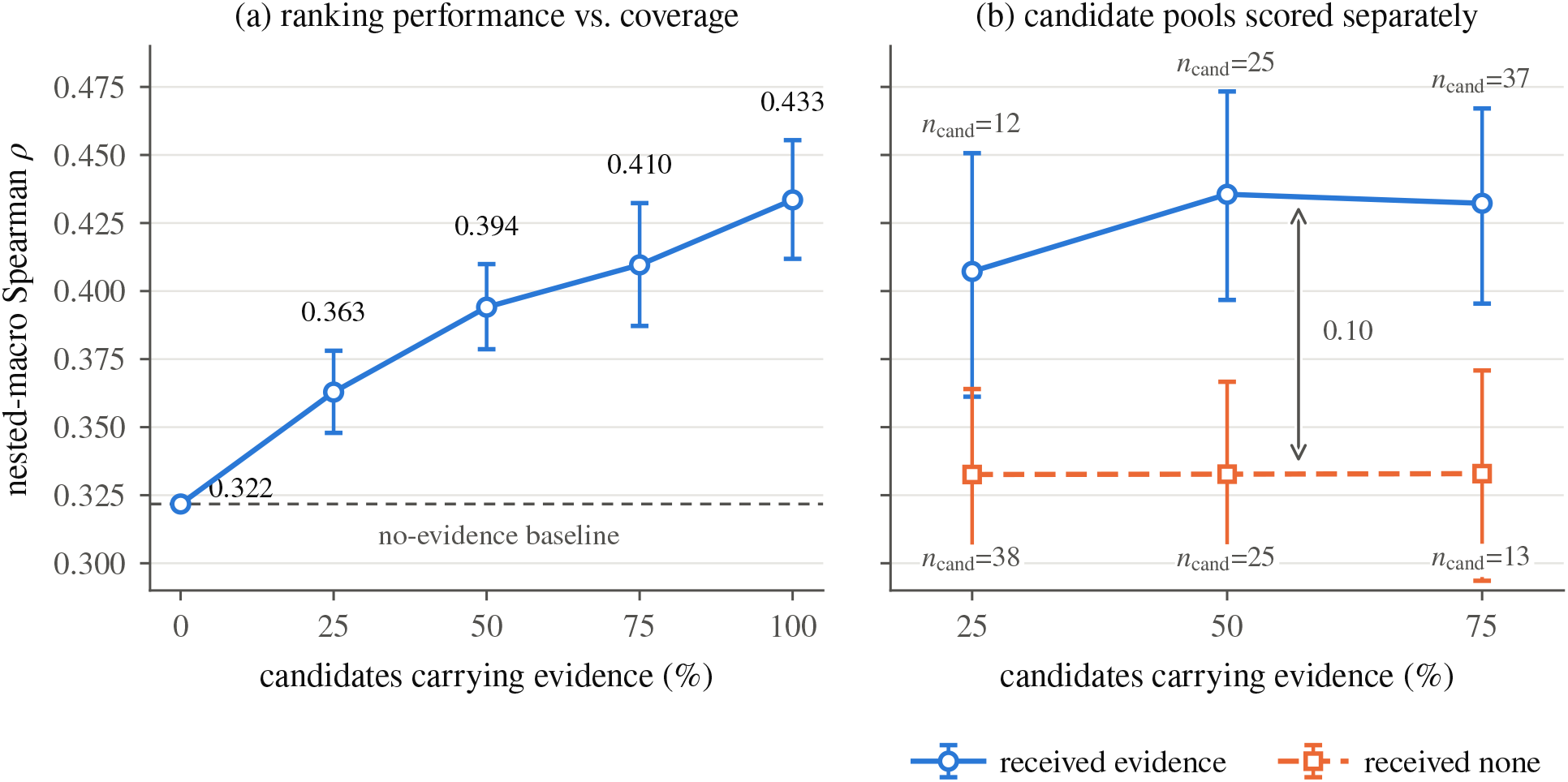
Evidence coverage and ranking performance. GPT-5.6 Sol at medium effort. **(a)** Score on 581 common episodes; intervals show paired differences from R, anchored at R. **(b)** Candidates with and without evidence, evaluated at each dose on its own episode coverage; intervals are for subgroup scores and *n*_cand_ gives subgroup size. All intervals use 10,000 protein-level bootstrap resamples.

#### Improvements are concentrated among candidates with evidence

At 25%, 50%, and 75% coverage, annotated candidates score 0.407–0.436, whereas unannotated candidates remain near 0.333 (Figure 2b). Gains therefore concentrate among annotated candidates. This exploratory split changes membership and subgroup size across doses, so it is not a controlled mixture decomposition.

### 5.2 Combining Structural and Evolutionary Evidence

We compare R, S, E, and SE for GPT-5.6 Sol at medium effort using the same prompt version. On the 581 episodes (195 assays, 166 proteins) scored by all four conditions, both modalities improve ranking individually, and combining them yields further gains over either modality alone (Table 2).

**Table 2:** Evidence modality ablation for GPT-5.6 Sol at medium effort on 581 common episodes (195 assays, 166 proteins), using the nested-macro aggregation in Section 4.2. Paired contrasts and intervals are reported below.

| Condition | $\rho$ | $\Delta$ vs. R |
| --- | --- | --- |
| R (no evidence) | 0.3218 | — |
| S (structure only) | 0.3986 | +0.077 |
| E (evolution only) | 0.4051 | +0.083 |
| SE (both) | 0.4335 | +0.112 |

SE exceeds S by 0.0349 (95% CI [0.0219, 0.0475]) and E by 0.0284 (95% CI [0.0135, 0.0428]). Intervals use 10,000 paired protein-level bootstrap resamples on this four-condition common panel (Section 4.2) and are not adjusted for multiple comparisons. These contrasts support an incremental benefit from combining modalities in the tested model and effort setting.

### 5.3 Comparison with Specialized Predictors

On each entry’s available coverage, GPT-5.6 Sol at medium effort moves from above 14 to above 65 of 95 published predictors under SE; Claude Opus 5 reaches 91 of 95 (Figure 3; Table 5). Claude Opus 5 and GPT-5.6 Sol at max effort also enter the *ρ≥* 0.45 range occupied by 17 specialists (Table 3). These descriptive positions are not paired tests, and weaker LLMs show much smaller gains relative to specialists.

**Table 3:** The *ρ≥* 0.45 band, our arms interleaved with the published predictors by own-coverage *ρ*. The matched column rescored all displayed entries on their common 254-episode panel (87 assays; 79 proteins); rows remain ordered by own coverage. Alignment/structure methods are marked A/S and sequence-only SEQ. Six further methods— RSALOR 0.466, ESM3 0.460, SaProt-650M-AF2 0.457, VespaG 0.455, GEMME 0.451, S2F 0.451—complete the band below the last row shown, so all three of our arms outrank them by own coverage. Arms marked *†*have not run the full 651 episodes.

| # | Entry | Own $\rho$ | Matched $\rho$ | Kind |
| --- | --- | --- | --- | --- |
| 1 | VenusREM | 0.523 | 0.535 | A/S |
| 2 | **GPT-6 Astra · max · SE$^\dagger$** | 0.514 | 0.520 | **ours** |
| 3 | ProSST-2048 | 0.512 | 0.524 | A/S |
| 4 | ProSST-4096 | 0.499 | 0.519 | A/S |
| 5 | S3F-MSA | 0.494 | 0.498 | A/S |
| 6 | **Claude Opus 5 · max · SE** | 0.486 | 0.492 | **ours** |
| 7 | S2F-MSA | 0.484 | 0.489 | A/S |
| 8 | ProSST-1024 | 0.483 | 0.509 | A/S |
| 9 | ESCOTT | 0.477 | 0.475 | A/S |
| 10 | ProSST-128 | 0.472 | 0.492 | A/S |
| 11 | PoET | 0.471 | 0.482 | A/S |
| 12 | S3F | 0.469 | 0.476 | A/S |
| 13 | ProSST-512 | 0.469 | 0.485 | A/S |
| 14 | **GPT-5.6 Sol · max · SE** | 0.466 | 0.480 | **ours** |

**Table 4:**
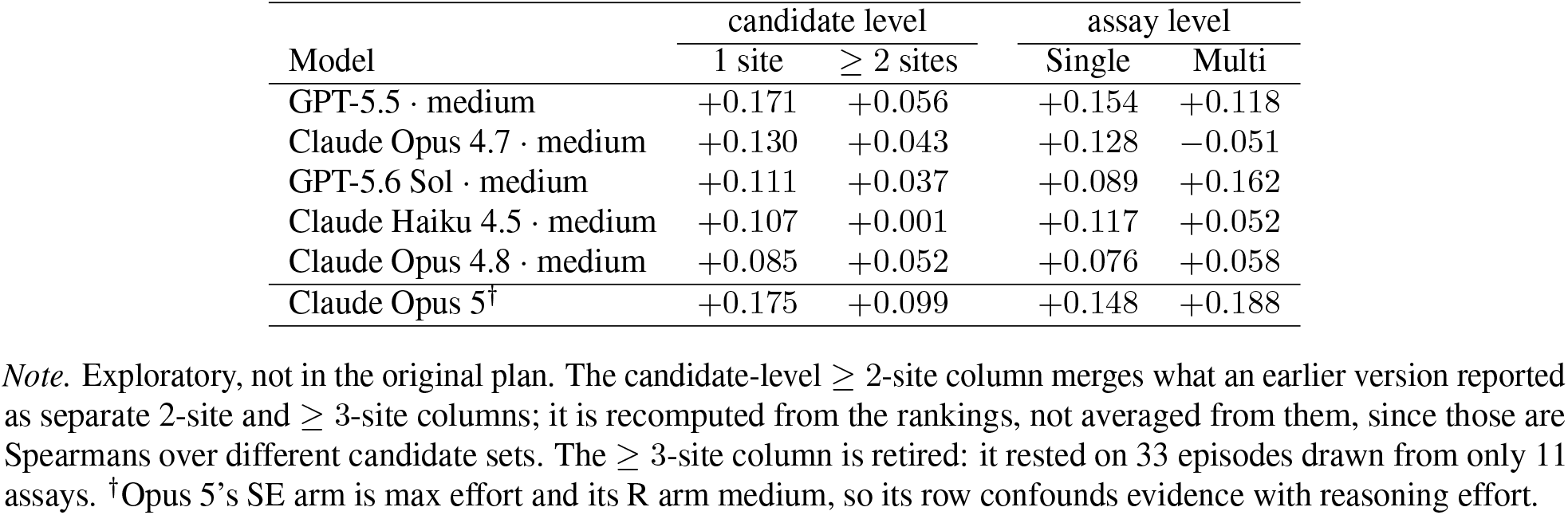
Evidence gain (SE−R) by mutation depth. *Candidate level* groups the candidates inside an episode by how many sites they mutate; *assay level* is the benchmark’s own Single/Multi label on the whole assay. The five models above the rule have a matched R/SE pair at the same effort; Opus 5^*†*^ is shown for reference only, as its pair crosses effort tiers, and it is excluded from Figure 6. The two levels disagree for GPT-5.6 Sol and for Opus 5. Columns are not comparable in magnitude—the groups differ in candidate count, fitness range and assay pool—only the SE−R difference within a column is. Depth strata are aggregated by a descriptive mean over assays rather than the five-category nested macro: that is the convention that reproduces the benchmark’s own published depth numbers, and inside a depth stratum the nested macro’s category layer is degenerate. Own coverage, point estimates. *Note*. Exploratory, not in the original plan. The candidate-level*≥* 2-site column merges what an earlier version reported as separate 2-site and *≥* 3-site columns; it is recomputed from the rankings, not averaged from them, since those are Spearmans over different candidate sets. The *≥*3-site column is retired: it rested on 33 episodes drawn from only 11 assays. ^*†*^Opus 5’s SE arm is max effort and its R arm medium, so its row confounds evidence with reasoning effort.

**Figure 3.**
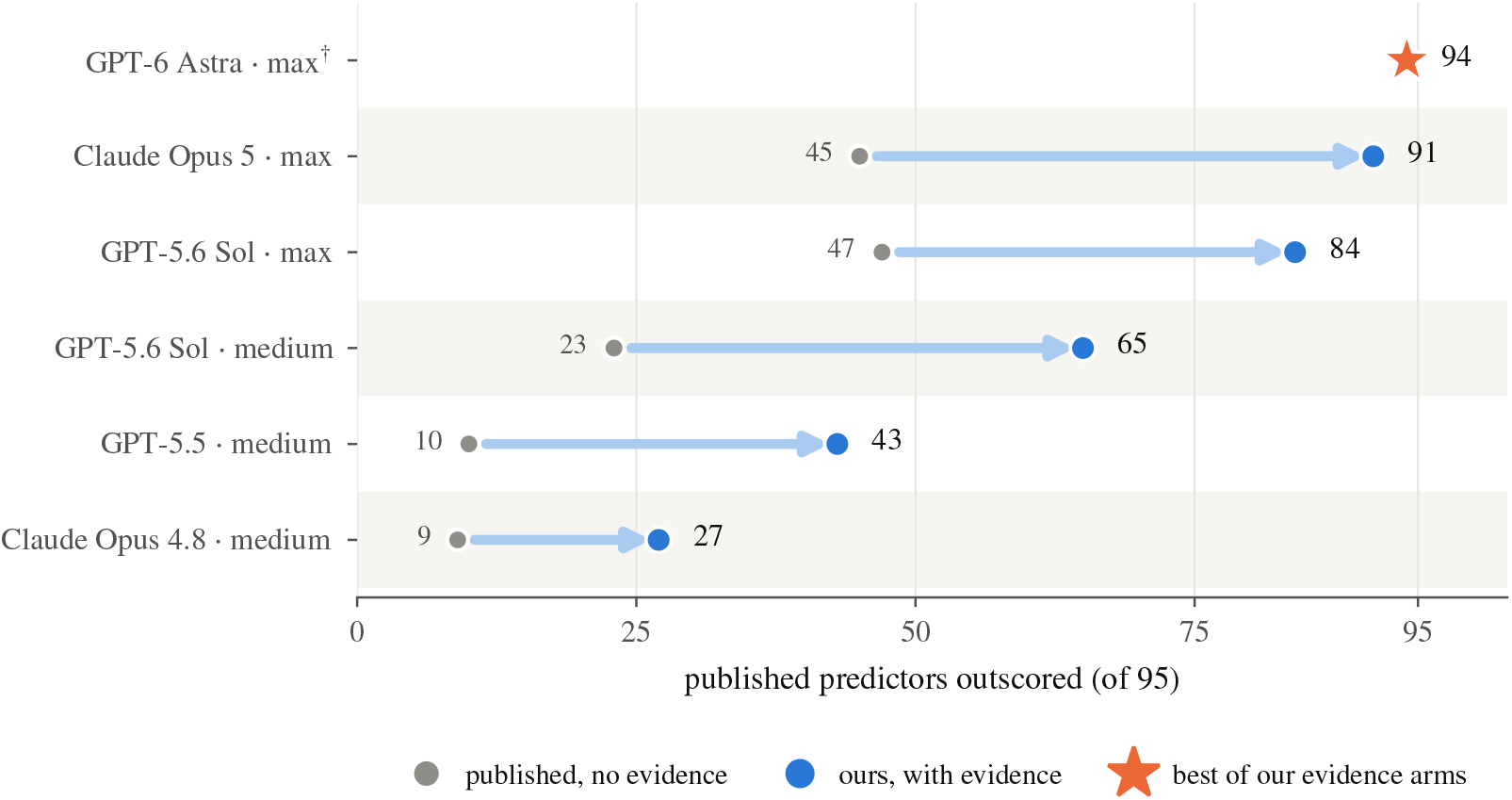
Descriptive leaderboard positions among 95 published predictors. Blue markers show our evidence conditions; grey markers show published no-evidence results for the same model and effort. Positions count predictors with lower own-coverage scores. Arrows combine evidence and channel differences and should not be interpreted as controlled evidence effects (Section 4.4). ^*†*^GPT-6 Astra has no published no-evidence counterpart and has scored only 267 of 651 episodes. Its incomplete, non-random coverage limits comparison with other entries.

#### Matched specialist comparisons are panel-dependent

On the strict 254-episode intersection of all 14 entries in Table 3, VenusREM, ProSST-2048, and GPT-6 Astra score 0.535, 0.524, and 0.520. GPT-6 differs by −0.015 from VenusREM (95% CI [−0.051, +0.019]) and −0.004 from ProSST-2048 ([−0.042, +0.031]). On the wider 561-episode pairwise panel, Claude Opus 5 remains 0.048 below VenusREM ([−0.081, −0.022]). GPT-6’s incomplete, non-random coverage limits generalization beyond its scored episodes.

### 5.4 Preliminary Evaluation on Post-Cutoff Assays

The three evidence conditions score 0.372–0.385 on their own coverage, above the strongest of 18 published no-evidence LLM baselines (0.347; Table 6). On 56 shared episodes, estimated advantages over that baseline are +0.043 for GPT-6 Astra (95% CI [0.013, 0.074]), +0.033 for GPT-5.6 Sol ([0.009, 0.058]), and +0.024 for Claude Opus 5 ([−0.021, 0.064]). These cross-channel comparisons do not isolate a within-model evidence effect.

VenusREM’s post-cutoff score is confounded by much shallower rebuilt alignments. In a 33-assay substitution control, its degradation exceeds that of GPT-5.6 Sol with evidence by 0.238 ([0.170, 0.315]), identifying alignment quality as a major factor (Appendix B.2). The 25-assay transfer result remains preliminary and does not isolate new-protein effects.

### 5.5 Evidence can substitute for additional LLM reasoning effort

For GPT-5.6 Sol, low effort with evidence exceeds medium effort without evidence by 0.104 (95% CI [0.084, 0.125]; 588 shared episodes); the corresponding Opus 4.8 contrast is not significant. Published max-effort comparisons cross channels (Figure 4). Because effort tiers are vendor settings, these results do not establish savings in latency, tokens, or cost.

**Figure 4.**
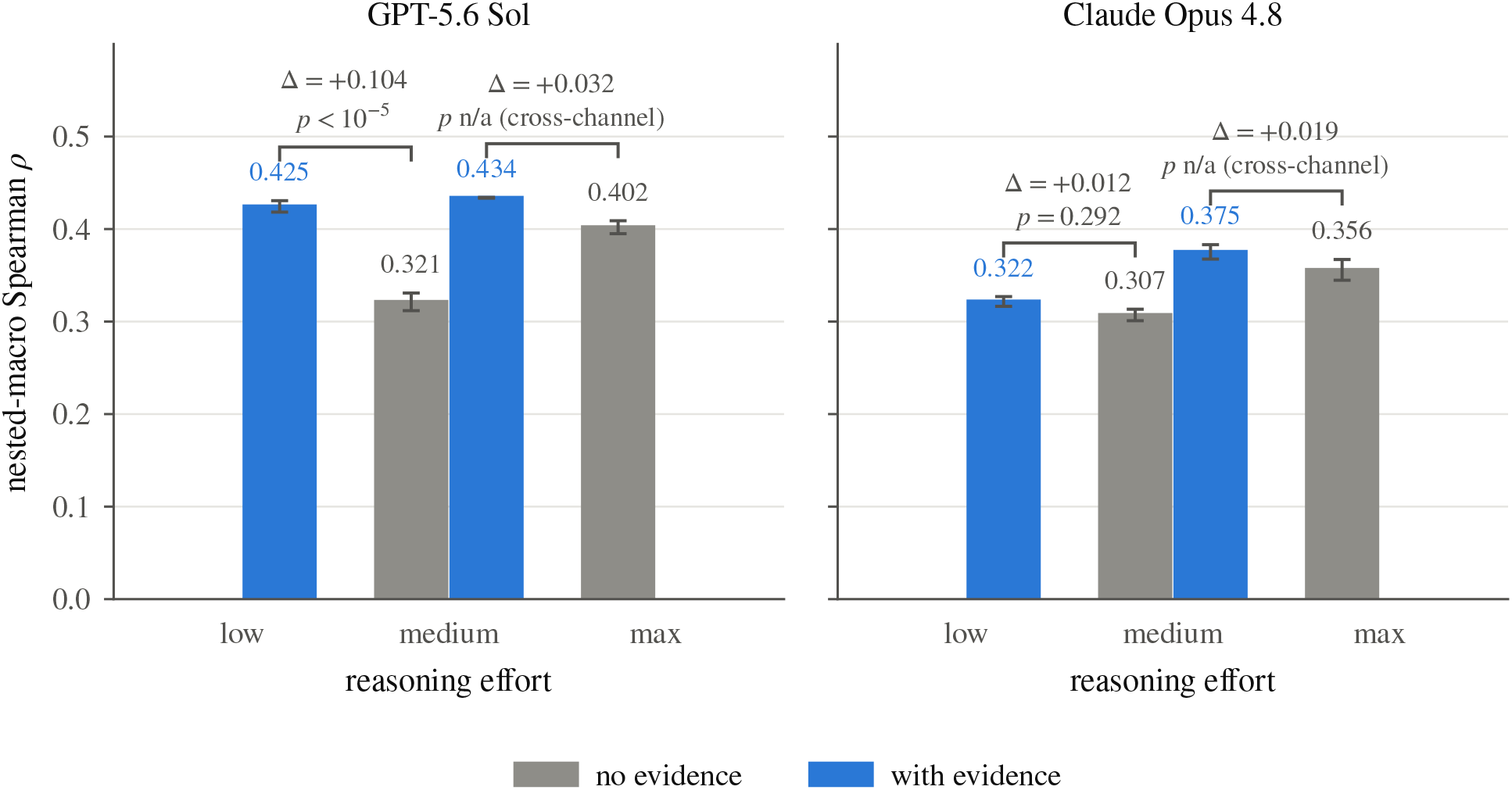
Evidence can substitute for additional reasoning effort. Bars show own-coverage scores; error bars are standard errors across draws. Same-channel brackets show paired differences and unadjusted protein-level permutation *p*-values. Published max-effort comparisons cross channels and have no *p*-values.

The substitution claim is specific to this model and effort pair; cross-version sensitivity results appear in Appendix B.3.

### 5.6 Performance beyond evolutionary rank fusion

An evidence-only readout does not test whether prompting beats a simple combination of evidence and the model’s no-evidence prediction. We therefore sum evolutionary log-frequency ratios over all mutated sites, then average the resulting candidate ranks with the no-evidence ranks at fixed *α* = 0.5. We score fusion on the same episodes and candidates as SE. Definitions, uncertainty, and sensitivity appear in Appendix C.

#### Fusion is a substantial comparison baseline

Fusion exceeds R for all six models. Relative to fusion, GPT-5.5 and GPT-5.6 Sol retain margins of +0.032 and +0.029, whereas Haiku 4.5, Opus 4.7, and Opus 4.8 fall below it by 0.094, 0.045, and 0.023; all five paired intervals exclude zero. Opus 5 retains a +0.036 margin ([+0.012, +0.057]) using public max-effort R, an effort-matched but cross-channel comparison. The signs persist against each model’s strongest tested weight. GPT-5.6 Sol also exceeds fusion on single-site candidates (+0.037, [+0.018, +0.055]; Appendix C).

Across models, the SE advantage over fusion is associated with no-evidence performance (Pearson +0.789; Spearman +0.829). The trend is not monotonic and vanishes for raw SE−R gains (Pearson −0.027; Spearman +0.086). This small, heterogeneous panel therefore supports an exploratory predictive association. Because the readout omits structure, it does not isolate integration ability or an internal reasoning mechanism.

#### Mutation depth

An exploratory analysis finds smaller evidence gains for multi-site variants under the descriptive aggregation, with some sensitivity to the estimator. Appendix B.4 reports the full analysis and its limitations.

## 6 Discussion

### Evidence access as an experimental axis

Performance rises with evidence coverage and falls when the same content is assigned to the wrong candidates. Evidence availability should therefore be studied alongside model and inference scaling, without implying a universal scaling law. Evidence construction and longer prompts also carry costs that must be measured before claiming efficiency gains.

### Implications for evaluating domain capability

LLM–specialist comparisons conflate parameters with input information. Evidence narrows the gap but does not establish equivalence: leading specialists retain small point-estimate advantages over GPT-6 on the strict panel, and VenusREM significantly exceeds Opus 5 on the wider pairwise panel. The temporal comparison is also sensitive to alignment quality. Mechanical fusion further shows that evidence prompting can underperform a simple rule: weaker models can underperform evolutionary rank fusion, whereas stronger models often exceed it. The readout uses all mutated sites but omits structure, so these margins do not isolate integration ability. Structure-aware and learned fusion on held-out proteins are natural follow-ups.

### Limitations and future work

Incomplete, non-random response coverage limits generalization beyond scored episodes; matched scoring does not remove this selection effect. Published baselines cross access channels, and some comparisons cross effort settings. The modality and coverage ablations use one model and effort; the descriptive SE−R mutation-depth summaries lack paired intervals. Evidence annotations also identify mutation positions, so the comparisons do not isolate biological information from mutation-localization assistance. Fusion correlations and the best-weight envelope are exploratory, and Opus 5 crosses channels. The temporal set has 25 assays; its 33-assay alignment control excludes proteins longer than 1,000 residues and, after its VenusREM result was known, one alignment whose evidence weighting was computationally prohibitive. Variants with three or more substitutions remain too sparse for separate analysis. Larger temporal panels, held-out fusion baselines, and end-to-end cost measurements are needed.

## 7 Conclusion

BioEvidence improves zero-shot protein ranking without parameter adaptation: gains scale with correctly assigned evidence, narrow the specialist gap, and can exceed those from additional reasoning effort. An exploratory six-model analysis associates stronger no-evidence performance with larger margins over evolutionary rank fusion. External evidence is therefore a complementary scaling axis for scientific prediction, while matched fusion remains essential for interpreting how models use it.

## AI use statement

LLMs were used both as experimental subjects and as assistants in preparing this work. Experimental outputs were collected through vendor CLI channels for GPT and Claude models. An LLM assistant also supported code development, analysis scripting, and manuscript drafting and editing. The authors are responsible for the experimental design, verification of analyses, and claims in the manuscript. Model rankings are scored deterministically using Spearman correlation against experimental fitness; no LLM judge is used for the reported ranking metric.

## Ethics statement

This study uses public benchmark data and involves no new wet-lab experiments or recruitment of human participants. ProteinGym aggregates previously published deep mutational scanning assays.

Protein fitness prediction has potential dual-use implications. Our evaluation is restricted to ranking fixed benchmark candidates using existing structural and evolutionary annotations, and introduces no new trained model weights. These restrictions define the scope of the study but do not eliminate risks associated with downstream applications of improved variant prediction.

Safety-related refusals are retained in the evaluation records and reported separately from ranking scores. Their frequency varies across models and input conditions; the vendor’s refusal labels do not by themselves determine whether a request is safe or unsafe. Appendix A reports the observed failure categories. Our statistical treatment of missing rankings is described in Section 4.4.

## Reproducibility statement

Our evaluation protocol and statistical procedures are described in Section 4, with prompt construction, coverage, and model-identity checks in Appendix A. Appendix E documents the frozen analysis artifacts, provenance, and random seeds, and Appendix C gives the mechanical fusion definition and full results. The supplementary code package provides analysis scripts and derived data; its README specifies which analyses run using the included files and which require external data or raw model records.

## A Additional Experimental Details

### A.1 Prompt structure

Each episode is presented as a single user turn. The R condition contains an assay header, the wild-type sequence, all 50 candidate sequences, and the ranking instruction. The example below abbreviates sequences with ellipses; the actual prompts contain complete sequences. Line wrapping is adjusted for display.

~~~
**Protein:** BRCA1_HUMAN (Homo sapiens)
**Assay (what is measured):** OrganismalFitness; Growth
**Higher experimental fitness = HIGHER value of the measured
property.**
**Wild-type sequence (1863 aa):**
MDLSALRVEEVQNVINAMQKILECPICLELIKEP…
**50 candidate mutant sequences to rank:**
M01: MDLSALRVEEVQNVINAMQKILECPICLELIKEP…
M02: …
…
M50: …
Rank all 50 mutants from the one you predict has the MOST
favorable
effect on the assayed property (highest fitness) to the LEAST.
Reason through the
ordering, then on the last line output ONLY the JSON object:
{“ranking”: [“M03”, “M27”, … all 50 ids, best to worst]}
~~~

The S, E, and SE conditions insert their evidence blocks between the candidate list and the ranking instruction, preserving the base task wording. Candidate identifiers `M01`–`M50` do not encode substitution identities.

### A.2 Arms, channels, and completeness

The primary snapshot contains 14 experiments: 12 are marked `final_complete`, and two (`exp _1_3a`, the specialist comparison, and `exp_1_3b`, the post-cutoff evaluation) are marked `provisional partial`. Completion refers to collection status and does not imply that every response yields a valid ranking.

Main-benchmark conditions target 651 episodes, while post-cutoff conditions target 75. GPT and Claude runs use `codex-cli` and `claude-code-cli`, respectively. The audit records run labels, model identifiers, effort settings where available, and prompt metadata. Claude Opus 5 SE combines two run labels covering 327 and 324 episodes. GPT-6 Astra SE combines a main-grid run with 240 records and a 20-assay subset run with 60 records, yielding scored coverage of 89 assays in the specialist comparison.

### A.3 Post-cutoff evidence construction

We rebuild alignments from full-length parent proteins with five Jackhmmer iterations against UniRef90 (June 2021), using a preregistered threshold of 0.5 bits per residue rather than ProteinGym’s per-assay threshold sweep (Notin et al., 2022). ProteinGym/EVE preprocessing removes sequences with more than 50% gaps, retains focus columns, and weights sequences with *θ* = 0.01 for viral proteins and 0.2 otherwise. Frequencies exclude gaps; entropy uses base-2 logarithms over the 20 standard amino acids. Structures come from AlphaFold DB v6 when a matching full-length model exists. Five remaining parents are folded with ColabFold 1.6.1 (five models, three recycles, no Amber relaxation), followed by the same DSSP extraction as in the main benchmark. The protocol records unavailable evidence rather than imputing or manually repairing it.

### A.4 Channel calibration

Our remote model runs use CLI channels, whereas the published benchmark uses native APIs. At matched model and effort settings, our GPT-5.6 Sol baseline scores 0.041 below the published baseline (95% CI [−0.064, −0.021]; 197 common assays), and GPT-5.5 scores 0.060 below it ([−0.085, −0.035]; 199 assays). For Claude Opus 4.8, the estimated difference is +0.012 ([−0.009, +0.032]; 185 assays). These results do not support a model-independent correction for channel effects. We therefore use in-house R conditions for controlled evidence comparisons and label comparisons with published LLM baselines as crossing channels. Differences in sampling settings, reasoning budgets, and refusal behavior remain possible confounders in those comparisons.

### A.5 Failure taxonomy

Refusals, context overflows, timeouts and unparsable responses are recorded by kind and excluded from ranking scores (Section 4.4). Their distributions differ substantially across models:

- **Safety blocks** dominate for the frontier arms. Out of 651 episodes: `opus5 max SE` 85, `opus48 med R` 64, `gpt56 med R` 62, `gpt55 med SE` 28. For the Claude channel these are further split by the vendor into `bio safeguard` and generic `other`; for `opus5 max SE`, 53 of 85 were `bio safeguard`.
- **Context overflow** reduces coverage for Claude Haiku 4.5. `haiku med R` fails on 1 episode but `haiku med SE` overflows on 38, consistent with the increased input length. Context failures contribute to the lower coverage of its SE condition.
- **Timeouts**. One timeout was recorded in `gpt56 max SE`.

Adding evidence does not move refusals in a consistent direction: it raises them for GPT-5.6 Sol at medium effort (62*→* 66) and lowers them for Claude Opus 4.8 (64 *→*38) and GPT-5.5 (50*→* 28). These counts are descriptive and do not identify a causal mechanism.

### A.6 Reference predictions

The comparison set comprises 95 published protein predictors and 43 site-hosted LLM entries. The primary snapshot lists episode-level source data for 13 main-benchmark predictors: ESM-1v (ensemble), ESM2-650M, EVE (ensemble), EVmutation, GEMME, MSA Transformer (ensemble), MULAN-small, ProGen2-xlarge, ProtSSN (ensemble), SaProt-650M-AF2, Site-Independent, Tranception-L, and VenusREM, at 651 episodes each except VenusREM at 642. Availability in the source data does not necessarily imply successful loading into every comparison; the provisional specialist experiment has the gaps listed below. Separately, the Table 3 matched analysis reconstructs all 11 displayed specialist rows directly from the official ProteinGym v1.3 variant-level prediction archive, with complete coverage of the 651 frozen episodes. The post-cutoff comparison draws on 20 reference arms including VespaG, ESM2-650M and 18 no-evidence LLM arms spanning GPT-5.5, GPT-5.6 Sol, Claude Opus 4.8 and Gemini 3.5 Flash at several effort tiers, and includes three in-house evidence conditions after the exclusion described in Appendix A.7.

### A.7 Served-model audit and excluded runs

A post-cutoff condition requested as Claude Opus 5.5 was excluded after response metadata revealed model substitution. Of 75 requests, 66 were answered by Claude Opus 5, eight were refused, and one was answered by Opus 5.5. Across the scored responses, Opus 5 accounted for 1,190,456 output tokens and Opus 5.5 for 361. A companion main-benchmark run showed the same issue and was stopped after 94 requests.

The original check tested whether the requested identifier appeared in response metadata. The revised audit instead measures the share of output tokens attributed to that model. Reported Claude conditions satisfy the *≥* 99.4% criterion.

A separate three-cell cross-model probe is also excluded from the reported experiments. GPT CLI records do not support an equivalent token-attribution check.

The excluded runs are retained for provenance and are not relabeled as Opus 5 conditions because their outputs mix models. Removing the post-cutoff condition changes the all-condition intersection from 54 to 56 episodes. Exclusion is based on model-identity validity, independently of its effect on statistical significance. Reported temporal results use the post-exclusion snapshot; the earlier snapshot is retained as an audit record.

### A.8 Remaining gaps in the provisional experiments

The primary specialist snapshot loaded episode-level scores for VenusREM (642) and GEMME (651) but omitted several other specialist rows. The supplemental ProteinGym v1.3 import closes this data gap for every specialist displayed in Table 3; it does not recover missing LLM responses. Consequently, 351 of 2604 expected in-house response records remain outstanding in `exp 1 3a`, and seven of 225 remain in `exp 1 3b`. These response-collection gaps underlie the provisional labels and remain distinct from specialist prediction availability.

## B Additional Results

### B.1 Extended specialist comparison

Table 5 complements the main-text comparison with predictor-family counts. It uses own coverage and remains subject to the coverage limitations in Section 5.3.

The own-coverage rankings complement the matched comparisons in Section 5.3; different response coverage prevents a direct paired interpretation of the leaderboard counts.

**Table 5:** Descriptive positions among 95 published predictors. Each entry uses its own coverage. Columns report the number of lower-scoring predictors overall and within the sequence-only and alignment/structure groups, followed by the difference from each group’s median (0.3745 and 0.4350). The published Opus 5 row reproduces the reference counts of 49/95, 41/46, and 8/49; it is a separate reference entry from the effort-specific comparisons in Figure 3.^*†*^GPT-6 Astra has incomplete run coverage. Other LLM entries also omit unsuccessful episodes. GPT-5.5 at medium effort with evidence (0.397, above 43 of 95) and Claude Opus 4.8 at medium effort with evidence (0.375, above 27 of 95) are omitted for brevity.

| Arm | $\rho$ | Scored | /95 | Seq /46 | A/S /49 | $\Delta$ seq | $\Delta$ A/S |
| --- | --- | --- | --- | --- | --- | --- | --- |
| GPT-6 Astra · max · SE<sup>†</sup> | 0.514 | 267/651 | 94 | 46 | 48 | +0.139 | +0.079 |
| Claude Opus 5 · max · SE | 0.486 | 561/651 | 91 | 46 | 45 | +0.111 | +0.051 |
| GPT-5.6 Sol · max · SE | 0.466 | 591/651 | 84 | 46 | 38 | +0.092 | +0.031 |
| GPT-5.6 Sol · medium · SE | 0.434 | 585/651 | 65 | 44 | 21 | +0.059 | −0.001 |
| GPT-5.6 Sol · low · SE | 0.425 | 590/651 | 60 | 44 | 16 | +0.050 | −0.011 |
| Claude Opus 5, published, no evidence | 0.406 | — | 49 | 41 | 8 | +0.032 | −0.029 |
| GPT-5.6 Sol · medium · R | 0.321 | 589/651 | 14 | 12 | 2 | −0.053 | −0.114 |

### B.2 Post-cutoff scores and coverage

Table 6 reports the descriptive scores discussed in Section 5.4, including the different coverage restrictions.

**Table 6:** Post-cutoff evaluation. Own-coverage scores use each entry’s available episodes (*n*). Matched scores use 56 common episodes, except VenusREM (^*\**^), which is evaluated on 55. The strongest of 18 published no-evidence LLM conditions is shown. Methods using evolutionary information receive alignments rebuilt for this study, which are substantially shallower than those of the main benchmark. Table entries are descriptive point estimates under the temporal study-group aggregation; paired study-group bootstrap intervals for all three comparisons with the no-evidence LLM baseline are reported in the text.

| Entry | | own cov. ($n$) | matched |
| --- | --- | --- | --- |
| GPT-6 Astra · max · SE | ours | 0.381 (65) | **0.373** |
| GPT-5.6 Sol · max · SE | ours | 0.385 (75) | **0.363** |
| Claude Opus 5 · max · SE | ours | 0.372 (66) | **0.353** |
| gpt-5.6-sol-max | published, no evidence | 0.347 (75) | 0.329 |
| VespaG | specialist | 0.310 (75) | 0.313 |
| VenusREM | specialist | 0.250 (74) | 0.225* |
| ESM2-650M | specialist | 0.175 (75) | 0.171 |

Rebuilt post-cutoff alignments have median effective depth 223, versus 5,102 for ProteinGym’s main-benchmark alignments. To measure this confound, we replace the official alignment with a post-cutoff-protocol alignment for 33 main-benchmark assays while holding proteins, variants, structures, and model settings fixed. VenusREM falls from 0.5263 to 0.2656 (−0.2607, 95% CI [−0.3284, −0.2022]), whereas GPT-5.6 Sol with evidence falls from 0.4429 to 0.4201 (−0.0229, [−0.0440, −0.0020]). The degradation difference is 0.238 ([0.170, 0.315]). This implicates alignment quality but does not isolate new-protein effects.

### B.3 Cross-version effort sensitivity

GPT-5.5 with evidence exceeds in-house GPT-5.6 Sol without evidence by 0.072 (95% CI [0.041, 0.102]; 588 shared episodes). Against the published GPT-5.6 Sol baseline, the estimate is 0.033 across 198 common assays ([0.00006, 0.06517]), but restricting to assays with all three draws scored yields 0.032 ([−0.001, 0.063]). This cross-channel comparison is therefore sensitive to coverage and does not define a general exchange rate between evidence, model version, and effort.

### B.4 Performance by Mutation Depth

Under the descriptive candidate-level aggregation, all five models with R and SE at the same effort show larger gains for single-site variants than for variants with two or more substitutions: GPT-5.5 +0.171 vs. +0.056, Opus 4.7 +0.130 vs. +0.043, GPT-5.6 Sol +0.111 vs. +0.037, Haiku 4.5 +0.107 vs. +0.001, Opus 4.8 +0.085 vs. +0.052 (Figure 6;full numbers, including the benchmark’s own assay-level split, in Table 4). The interface supplies site-level annotations and does not explicitly model interactions between substitutions, which may limit its utility for multi-site variants.

This pattern depends partly on aggregation. For Opus 4.8, the ordering changes under alternative estimators: under the paired matched-coverage estimator it reverses to +0.068 vs. +0.087, and under the headline nested macro it flattens to +0.075 vs. +0.080. The other four keep their ordering under all three conventions.

The split is per candidate rather than by the benchmark’s own per-assay Single/Multi label, because 65 of its 69 Multi-mutant assays mix depths—a median 51% of their candidates mutate a single site—so the assay label says little about the candidates inside it. Table 4 reports both.

## C Mechanical Fusion: Definition and Full Results

### Evidence readout and ranking conventions

The deterministic score sums over the mutated positions 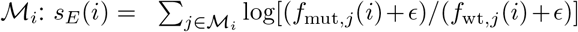, with *ϵ* = 10^−9^. Frequencies are read from every mutated site’s evolutionary block in the actual SE prompt; parsed positions are checked against the complete mutation list. No weights are fitted to fitness labels. This readout uses evolutionary frequencies but no structural features. Candidate identifiers break ties when converting *s*_*E*_ into the ordering *r*_*E*_. Let *r*_*R*_(*i*) denote the no-evidence rank. Candidate-level fusion is

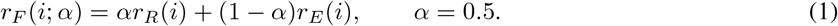

Spearman correlation is then computed from −*r*_*F*_ and measured fitness, assigning average ranks to tied fusion scores. Thus equal fusion scores are not arbitrarily ordered by identifier. This ranking convention differs from directly correlating the scalar evidence scores with fitness, so the rank-based evidence scores here need not exactly match a direct-score readout.

### Coverage and paired uncertainty

All 651 SE prompts yield evidence scores for all 50 candidates. On each model’s R/SE intersection, R, *E*_ro_, F, and SE are evaluated on the same complete candidate lists; both the minimum and median candidate count are 50. The common episode sets differ across models and are listed in Table 7. For SE−F at fixed *α* = 0.5, intervals are percentile 95% paired protein-level bootstrap intervals with 10,000 resamples, using the shared analysis seed `20260922`. They are not adjusted for the six model comparisons. Five pairs use in-house R; Opus 5 uses public max-effort R and therefore crosses access channels. The public PG-LLM Opus 5 entry supplies 541 max-effort R rankings. Each was mapped from its mutation labels to the frozen local candidate identifiers and verified as an exact 50-candidate set; 532 also have a scorable in-house max-effort SE response. The public and locally recomputed episode Spearman values agree to within 5.0*×* 10^−5^, the precision of the published values. This public-R analysis is same-effort but cross-channel.

**Table 7:** Matched component and fusion scores. *n* is the number of common episodes. *E*_ro_ is the deterministic readout, not an LLM given only evolutionary evidence. Δ_best_ is *ρ*_SE_−max(*ρ*_*R*_, *ρ*_*E*_); Δ_*F*_ is *ρ*_SE_−*ρ*_*F*_ at fixed *α* = 0.5. Intervals apply to Δ_*F*_. Opus 5 uses public max-effort R; the other R components are in-house.

| Model | $n$ | R | $E_{ro}$ | F | SE | $\Delta_{best}$ | $\Delta_F$ | 95% CI |
| --- | --- | --- | --- | --- | --- | --- | --- | --- |
| Haiku 4.5 | 609 | 0.152 | 0.345 | 0.320 | 0.226 | −0.119 | −0.094 | [−0.120, −0.070] |
| Opus 4.7 | 592 | 0.211 | 0.348 | 0.344 | 0.299 | −0.049 | −0.045 | [−0.064, −0.026] |
| GPT-5.5 | 597 | 0.245 | 0.344 | 0.360 | 0.392 | +0.048 | +0.032 | [+0.013, +0.050] |
| Opus 4.8 | 570 | 0.311 | 0.349 | 0.397 | 0.375 | +0.025 | −0.023 | [−0.040, −0.006] |
| GPT-5.6 Sol | 582 | 0.322 | 0.348 | 0.405 | 0.434 | +0.086 | +0.029 | [+0.011, +0.048] |
| Opus 5 (public R) | 532 | 0.406 | 0.353 | 0.454 | 0.490 | +0.084 | +0.036 | [+0.012, +0.057] |

On this triple-shared coverage, public max-effort R exceeds in-house medium-effort R by 0.061 (95% CI [+0.039, +0.085]), and its equal-weight fusion exceeds the medium-R fusion by 0.031 ([+0.015, +0.047]). Thus the weaker medium-effort component overstates the descriptive Opus 5 fusion margin, but the margin is not eliminated by the stronger same-effort component. For the public-R row, fusion scores at *α* = 0.25, 0.5, 0.75 are 0.412, 0.454, and 0.444; SE exceeds the strongest by 0.036. For the in-house medium-R row on these same episodes, the corresponding scores are 0.401, 0.424, and 0.392.

### Fixed weights and the post-hoc envelope

The primary comparison fixes *α* = 0.5 and fits no weights to fitness labels. The alternatives *α*∈ {0.25, 0.75} are sensitivity checks, with lower *α* assigning more weight to evidence. We also summarize the strongest fusion score among these three weights separately for each model. Choosing this weight uses its benchmark fitness correlations, so the resulting envelope is a post-hoc, benchmark-informed comparison, not a held-out estimate of a tuned predictor. No confidence intervals or significance claims are assigned to the selected-weight margins. The correlations with R performance are exploratory summaries of six model points, with different common episode sets and an access-channel mismatch for Opus 5.

### All-site readout and candidate-depth sensitivity

For GPT-5.6 Sol, replacing a first-site-only readout with the all-site sum raises fusion from 0.3847 to 0.4046 on the same 582 episodes (+0.0199, 95% CI [0.0126, 0.0278]). The SE−F margin decreases from 0.0493 to 0.0294 (95% CI [0.0105, 0.0475]). We additionally restrict each episode to candidates with exactly one or at least two substitutions, rerank both components within that subset, and recompute fixed-weight fusion and Spearman correlation. Subsets with fewer than three candidates are excluded jointly. These are restricted-list analyses of existing responses, not new LLM calls. Table 10 uses the same nested-macro estimator and protein-level bootstrap as the main fusion analysis; subgroup intervals are unadjusted. The single-site margin remains positive, while the multi-site interval includes zero. First-site and all-site rules coincide exactly on single-site candidates. These controls address omitted mutated sites, but not the readout’s omission of structure.

**Table 8:** Opus 5 R-source sensitivity on identical coverage. Both rows use the same 532 episodes, all 50 candidates per episode, the same deterministic readout, and the same in-house max-effort SE rankings. The public max-effort R row matches effort but crosses access channels; the in-house medium-effort R row matches channel but crosses effort tiers. Intervals are paired protein-level bootstrap intervals for SE−F at *α* = 0.5.

| R source | $n$ | R | $E_{ro}$ | F | SE | SE–F | 95% CI |
| --- | --- | --- | --- | --- | --- | --- | --- |
| Public max effort | 532 | 0.406 | 0.353 | 0.454 | 0.490 | +0.036 | [+0.012, +0.057] |
| In-house medium effort | 532 | 0.345 | 0.353 | 0.424 | 0.490 | +0.066 | [+0.034, +0.095] |

**Table 9:** Fusion-weight sensitivity. Columns show fusion scores for the three fixed weights; *α*^*\**^ maximizes each model’s aggregate fusion score over this set. The final column is the descriptive margin against that post-hoc envelope. Scores are reported to four decimal places. Opus 5 uses public max-effort R and crosses access channels.

| Model | $\rho_{F,0.25}$ | $\rho_{F,0.5}$ | $\rho_{F,0.75}$ | $\alpha^*$ | SE—best fusion |
| --- | --- | --- | --- | --- | --- |
| Haiku 4.5 | 0.3558 | 0.3200 | 0.2301 | 0.25 | −0.1297 |
| Opus 4.7 | 0.3637 | 0.3438 | 0.2761 | 0.25 | −0.0648 |
| GPT-5.5 | 0.3692 | 0.3604 | 0.3027 | 0.25 | +0.0228 |
| Opus 4.8 | 0.3866 | 0.3974 | 0.3586 | 0.5 | −0.0228 |
| GPT-5.6 Sol | 0.3897 | 0.4046 | 0.3704 | 0.5 | +0.0294 |
| Opus 5 (public R) | 0.4123 | 0.4542 | 0.4436 | 0.5 | +0.0356 |

**Table 10:** GPT-5.6 Sol fusion by candidate mutation depth. All-site readout, *α* = 0.5, medium effort. Counts are episodes/assays/proteins. Candidate counts include repeated appearances across draws. Intervals apply to SE−F.

| Candidates | Counts | F | SE | SE−F | 95% CI |
| --- | --- | --- | --- | --- | --- |
| All | 582/195/166 | 0.4046 | 0.4340 | +0.0294 | [+0.0105, +0.0475] |
| Single-site | 539/183/156 | 0.3912 | 0.4280 | +0.0368 | [+0.0177, +0.0554] |
| Multi-site | 204/68/65 | 0.4067 | 0.4475 | +0.0409 | [−0.0105, +0.0755] |

All, single-site, and multi-site panels contain 29,100, 23,146, and 5,929 candidate appearances, respectively (27,828, 22,001, and 5,802 unique assay–variant pairs). An episode may contribute to both depth subsets.

## D Extended Related Work

Sequence models such as ESM-1v (Meier et al., 2021) and ESM2 (Lin et al., 2023) build on unsupervised protein representation learning (Rives et al., 2021). Alignment-based methods include EVmutation (Hopf et al., 2017), DeepSequence (Riesselman et al., 2018), EVE (Frazer et al., 2021), GEMME (Laine et al., 2019), MSA Transformer (Rao et al., 2021), and PoET (Truong & Bepler, 2023); Tranception (Notin et al., 2022) combines sequence modeling with retrieval. VespaG (Marquet et al., 2024) uses sequence embeddings at inference time but is trained on GEMME-derived labels. Structure-aware predictors include ESM-IF (Hsu et al., 2022), SaProt (Su et al., 2024), ProSST (Li et al., 2024), and S3F (Zhang et al., 2024); SaProt uses the structural alphabet introduced by Foldseek (van Kempen et al., 2024). VenusREM (Tan et al., 2025) combines evolutionary and structural context and is the highest-scoring published predictor in our main-benchmark comparison. EvoIF (Jiao et al., 2026) and RSALOR (Tsishyn et al., 2025) further motivate studying compact evolutionary and structural summaries. We ask how effectively a frozen general-purpose model can use related annotations through a text interface. We exclude supervised fitness predictors such as ProteinNPT (Notin et al., 2023b) and Kermut (Groth et al., 2024), and clinical pathogenicity predictors such as AlphaMissense (Cheng et al., 2023), whose supervision or prediction targets differ from our zero-shot ranking setting.

## E Reproducibility Details

### Results and provenance

The primary analyses use `paper_v2_analysis_snap10`, which recomputes paired intervals with category-preserving protein resampling on the frozen response set. Its `snapshot_manifest.json` records input response paths, modification times, and file sizes. Experiment status and coverage are recorded in `paper_experiment_status.csv`. The primary specialist collection and post-cutoff comparison remain labeled `provisional_partial`. Supplementary analyses use separate artifacts. The strict matched column in Table 3 is stored in `table3_matched_v13`; `score_table3_matched.py` restricts the official ProteinGym v1.3 variant-level predictions to the frozen candidate draws and intersects them with the three LLM arms, yielding 254 common episodes. All 11 displayed specialists have complete 651-episode coverage before that intersection. The three specialist contrasts quoted in the text use corrected intervals stored in `table3_matched_v13_ci_snap10`, recomputed from the frozen per-episode scores with 10,000 protein re-samples (seed 20260925). The modality results in Table 2 use the frozen snap9 four-arm intersection (581 episodes, 195 assays, 166 proteins), aggregated with the same nested-macro estimator as the primary analyses. The script `pv2_modality_paired ci.py` verifies all 2,324 cached scores against the raw rankings and computes the paired intervals with seed `20260922`. Point estimates, paired contrasts, and an input audit are stored in `paper_v2_modality_paired_ci_snap9_20260926`. Fusion scores, weight sensitivity, and summary statistics for the six-model panel are stored in `paper_v2_fusion_all_sites_panel_snap10_20260926`. This reporting subset uses five in-house R components and the public max-effort R component for Opus 5, then recomputes the cross-model correlations. Figure 5 is generated from these CSV files. The script `pv2_fusion_all_sites_panel.py` validates every mutated position, reproduces the original first-site panel, and recomputes all-site fusion and the Opus 5 R-source sensitivity on frozen responses. The candidate-depth analysis uses `pv2_fusion_all_sites.py`, with outputs in `paper_v2_fusion_all_sites_gpt56_snap10_20260926`. Public traces were pinned to site version `20260902gemini38`; 541 rankings passed exact mutation-set binding against the frozen local episodes, and neither public reasoning text nor public fitness values were retained. The fusion comparison uses the 532 episodes shared with the in-house max-effort SE arm. The dose-subgroup plot uses `paper_v2_dose_own_split_snap10`. The candidate- and assay-level depth analyses use `paper_v2_depth_merge_snap10` and `paper_v2 _depth_strata_snap10`; their descriptive point estimates are unchanged by the bootstrap correction. The corrected temporal common panels and study-group bootstrap intervals are stored in `paper_v2_common_panel_snap10`; they use seed `20260922`. The frozen alignment and structure-generation settings are recorded in `p5_protocol_frozen.json`. The alignment-substitution control is stored in `paper_v2_msa_control_33` and `paper_v2_msactl_llm`; both use protein-level bootstrap seed `20260925` and the same 33-assay panel.

**Figure 5.**
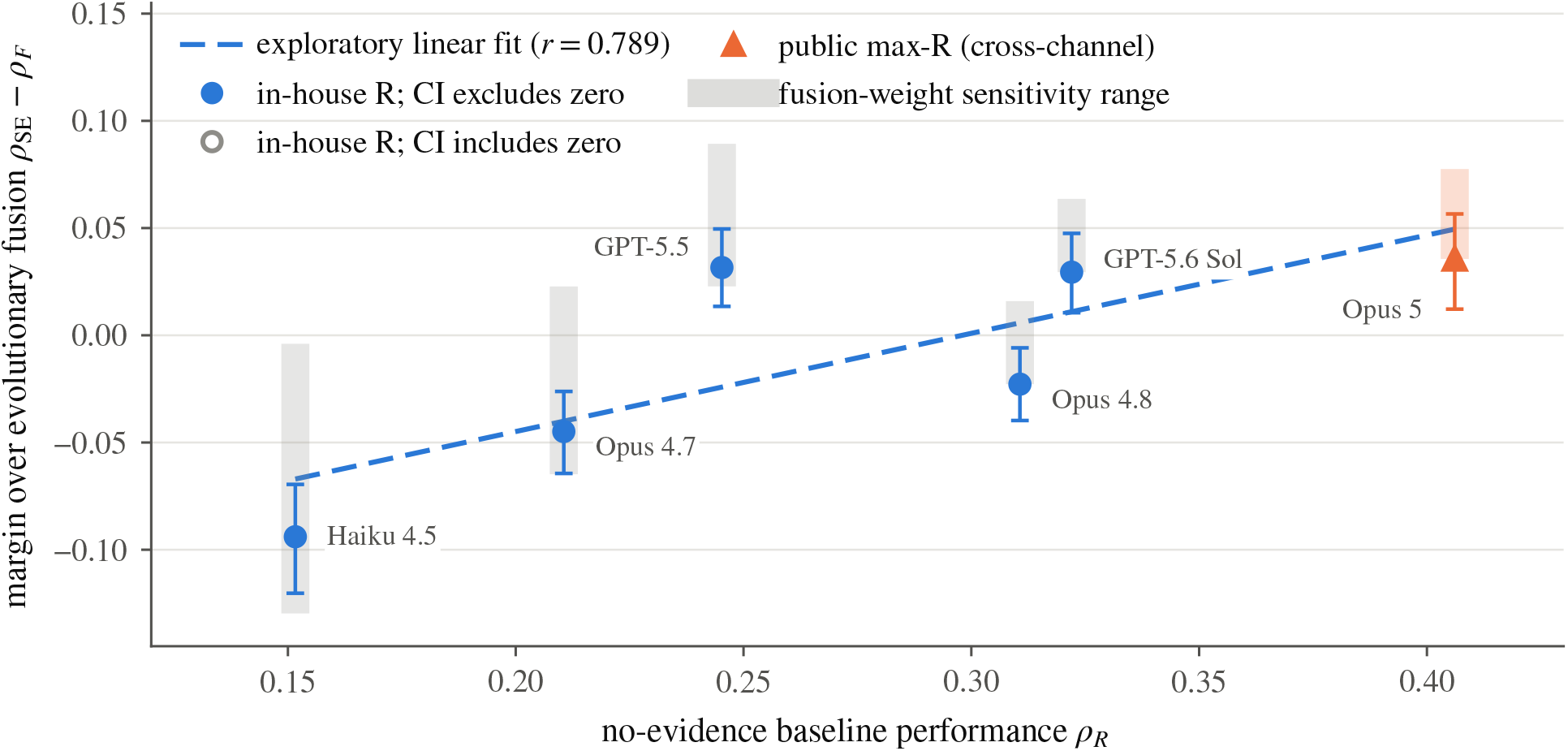
Performance beyond evolutionary fusion is associated with baseline model performance. Points show SE− fusion at fixed *α* = 0.5; the line is an exploratory six-model fit (Pearson +0.789). The readout sums log-ratios over all mutated sites. Error bars are paired bootstrap 95% intervals; shaded ranges span three tested weights. Opus 5 uses public max-effort R and crosses channels. Full results are in Appendix C.

**Figure 6.**
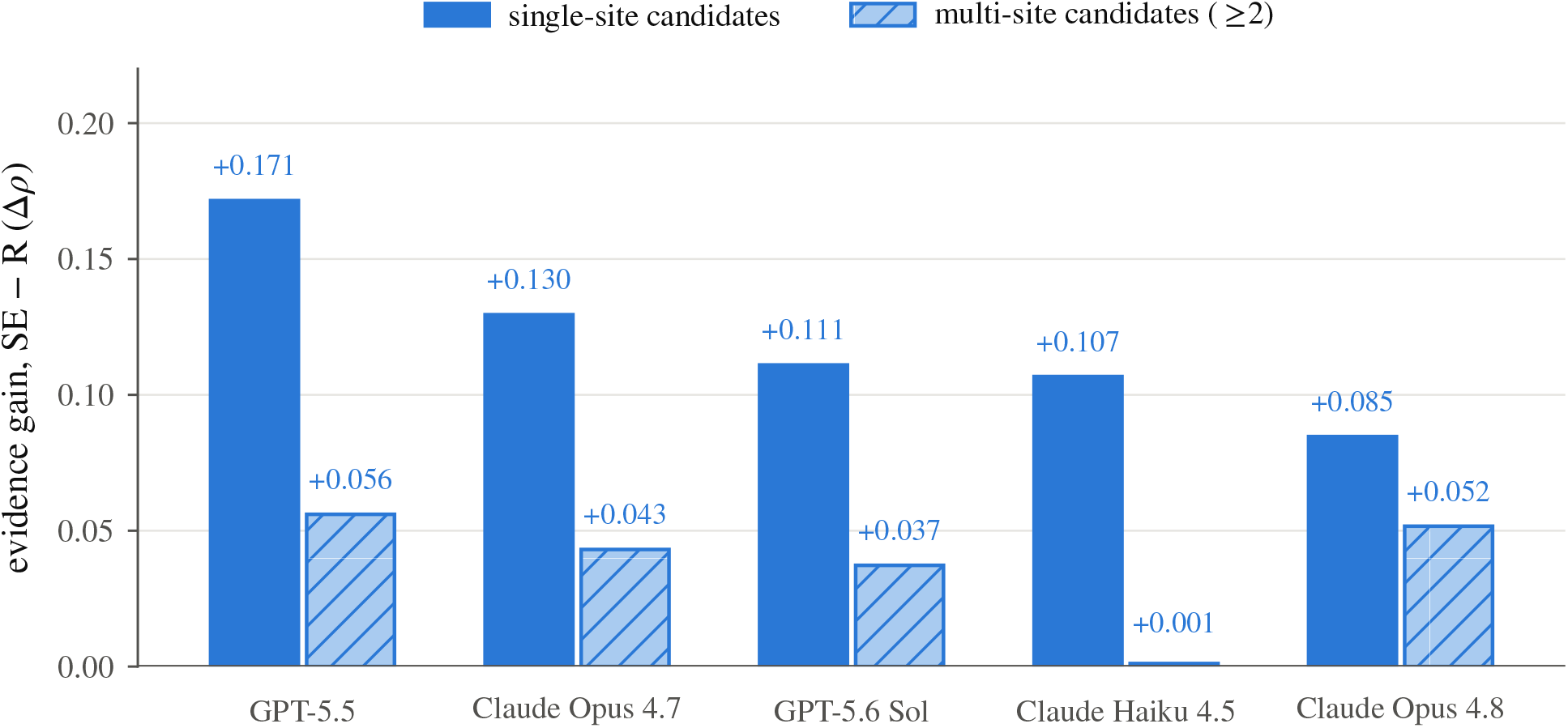
Evidence gain by mutation depth under descriptive aggregation. For each of the five models with a matched R/SE pair at the same reasoning effort, candidates inside an episode are grouped by how many sites they mutate and ranked against their own measured fitness, using the model’s original ranking restricted to that group. Bars give SE −R within each group, on own coverage; models are ordered by their single-site gain. Opus 5 is absent because it has no medium SE arm, so its only R/SE pair crosses effort tiers.

### Fixed candidates and prompts

We retain the benchmark’s fixed candidate draws: 217 assays, three draws per assay, and 50 candidates per draw, giving 651 episodes. The snapshot audit reports no duplicate cells or response directories outside the episode list. Remote model records identify the prompt version as `ranking-v1`. Claude effort settings are documented in job configurations rather than the response effort field, limiting uniform metadata verification across channels.

### Model identity and statistics

For Claude runs, the served-model audit measures the requested model’s share of output tokens; reported conditions meet the ≥99.4% criterion. Runs that failed this check are excluded (Appendix A.7). The GPT records do not expose an equivalent field. Primary confidence intervals use 10,000 paired protein-level bootstrap resamples; analysis seeds are specified in the corresponding scripts. The shared paper-analysis implementation uses seed `20260922`; supplementary scripts may use different fixed seeds.

### Reproducibility scope

Stored responses, candidate definitions, prompts, and scoring code support reproduction of the analyses without querying a model again. Fresh model runs need not reproduce the stored outputs because sampling and hosted model versions can change.

